# Number of ancestors and length of identity-by-descent tracks over time

**DOI:** 10.1101/2024.08.28.610091

**Authors:** Mikkel Spallou Eriksen, Gabriel Renaud

**Affiliations:** Department of Health Technology, Technical University of Denmark, Kongens Lyngby, Denmark

## Abstract

The divergence between genetic ancestors (those who have contributed to an individual’s DNA) and genealogical ancestors (individuals recognized as ancestors in a societal context) evolves significantly over time. This study focuses on small, insular populations, where the number of genetic ancestors can swiftly encompass the entirety of genealogical ancestors. Additionally, in these populations, runs of homozygosity (ROHs) frequently occur due to random chance. This study used a forward-in-time simulation based on the Wright-Fisher model to analyze genealogical and genetic relatedness, haplotype block lengths, and runs of homozygosity (ROH) across generations. In small, insular populations, the number of genetic ancestors rapidly encompassed the total number of potential ancestors, and ROHs occurred frequently due to genetic drift. Our results showed that while genealogical ancestors followed an initial exponential growth phase before stabilizing, genetic ancestors grew more slowly. This was especially the case when the initial population size was large. These findings highlight the high frequency of ROHs in smaller populations, providing insights into the genetic structure of historically isolated groups.

## Introduction

The study of population and evolutionary genetics is centered on understanding the genetic composition of populations and the mechanisms that drive genetic change over time. A population, defined as a group of interbreeding individuals of the same species within a specific geographical area, maintains a shared gene pool (1). This gene pool, which includes all alleles present in the population, allows for the calculation of allele frequencies, offering insights into genetic diversity and its role in adaptation and evolution (2). Genetic diversity is critical for a population’s resilience to environmental challenges, such as diseases, by increasing the likelihood that some individuals possess resistance or immunity, thereby ensuring the survival and reproduction of the population (3). The effects of insularity on genetic diversity have been a focus of various research efforts (4).

An illustrative example of an insular population is the Faroe Islands, a small, isolated community of approximately 50,000 individuals with a notable prevalence of rare diseases, such as Primary Carnitine Deficiency (5; 6). The Faroe Islands have a unique genetic history, with the population initially settling between the 4th and 6th centuries AD (7). The community remained small for centuries before experiencing growth in the 1800s (8). Gislason’s study (6) provided valuable insights into the genetic landscape and historical ancestry of this population. By utilizing whole genome sequencing of eleven individuals, the study revealed close relatedness among the population and identified a bottleneck event between the 1st and 4th centuries AD. Ancestry analysis indicated European and Admixed American origins, with the Faroese clustering near central European and British populations—a result of prolonged isolation and genetic drift. Despite their European roots, the Faroese exhibit levels of inbreeding comparable to those found in ancient European populations (6).

The genetic variance observed in populations is shaped by several evolutionary forces: genetic drift, gene flow, natural selection, and mutation (2). Genetic drift, particularly pronounced in smaller populations, refers to changes in allele frequency across generations due to chance events, leading to decreased genetic variation over time. This reduction can result from phenomena like the bottleneck effect or the founder effect, where significant reductions in population size or the establishment of new populations by a small group of individuals can lead to reduced genetic diversity (2).

At the molecular level, genetic variance is further influenced by recombination and independent assortment during meiosis. Recombination, occurring during Prophase I of meiosis, leads to the exchange of genetic material between maternal and paternal chromosomes, while independent assortment ensures that offspring inherit a varied combination of genes from both grandparents (9; 10; 2). This molecular variance contributes to the overall genetic diversity within a population.

Identity by descent (IBD) refers to identical DNA segments inherited from a common ancestor without recombination, and the length of these segments helps to study heredity and relatedness in populations (11). In small populations, runs of homozygosity (ROH), which reflect the effects of inbreeding, are common due to the limited number of distinct ancestors, underscoring the impact of isolation and population bottlenecks (12). This phenomenon was also observed in Gislason’s study of the Faroese population (6).

The Wright-Fisher model is a key tool in simulating genetic drift in populations over time, operating under several idealized assumptions to simplify the process (13; 14). This model allows for the simulation of genetic dynamics across generations, illustrating how some alleles become fixed while others are lost, particularly in small populations.

As one traces their ancestry back in time, they encounter two types of ancestors: genealogical ancestors, who are part of the family pedigree, and genetic ancestors, who have actually contributed genetic material. While these groups often overlap in recent generations, they can diverge further back in time due to the stochastic nature of meiosis, where only half of a person’s genes are passed on to their offspring.

In this study, we aim to quantify the number of genealogical and genetic relatives within a small isolated population through simulations. Building on the work of Coop (15; 16), who developed a formula to estimate the expected number of genetic ancestors in ancestral generations, we apply these methods to understand the rate at which runs of homozygosity (ROHs) increase over time in such populations. Coop’s formula predicts the number of genetic ancestors by considering the number of recombination events per transmitted genome, providing a framework for our investigation into the genetic dynamics of small, isolated populations. Our simulations written in Python can help studies looking at genealogical versus genetic ancestry.

## Methods

We implemented a forward-in-time simulation based on the Wright-Fisher model, capable of tracking gene spread, recombination effects, and the inheritance of genetic material across generations. The model assumes discrete-time generations, constant population size, random mating, and equal probability of recombination on every autosomal locus. The model uses parameters such as population size, number of generations, recombination rate, and sequence lengths of each chromosome. The model computes quantitative data, including average number of ancestors and descendants per generation, length and count of haplotype blocks, and length and frequency of runs of homozygosity (ROH). The code is available on https://github.com/mikkel-eriksen/ibd-over-time-public (17).

### Population initialization

To initialize the population, the model adopts a generation-by-generation approach. In each generation, excluding the initial one, individuals are assigned random parents from the preceding generation. The population structure resembles a Wright-Fisher model, but with the unique feature of each individual having two parents (Figure 1), similar to the method presented by Chang (18). The entire genome of an individual is represented as two homologous sequences (maternal and paternal), with chromosomes inherited from one parent concatenated into a single sequence. Break positions, representing both recombination events and the mechanism of independent assortment, are meticulously assigned for each parent.

**Figure 1:**
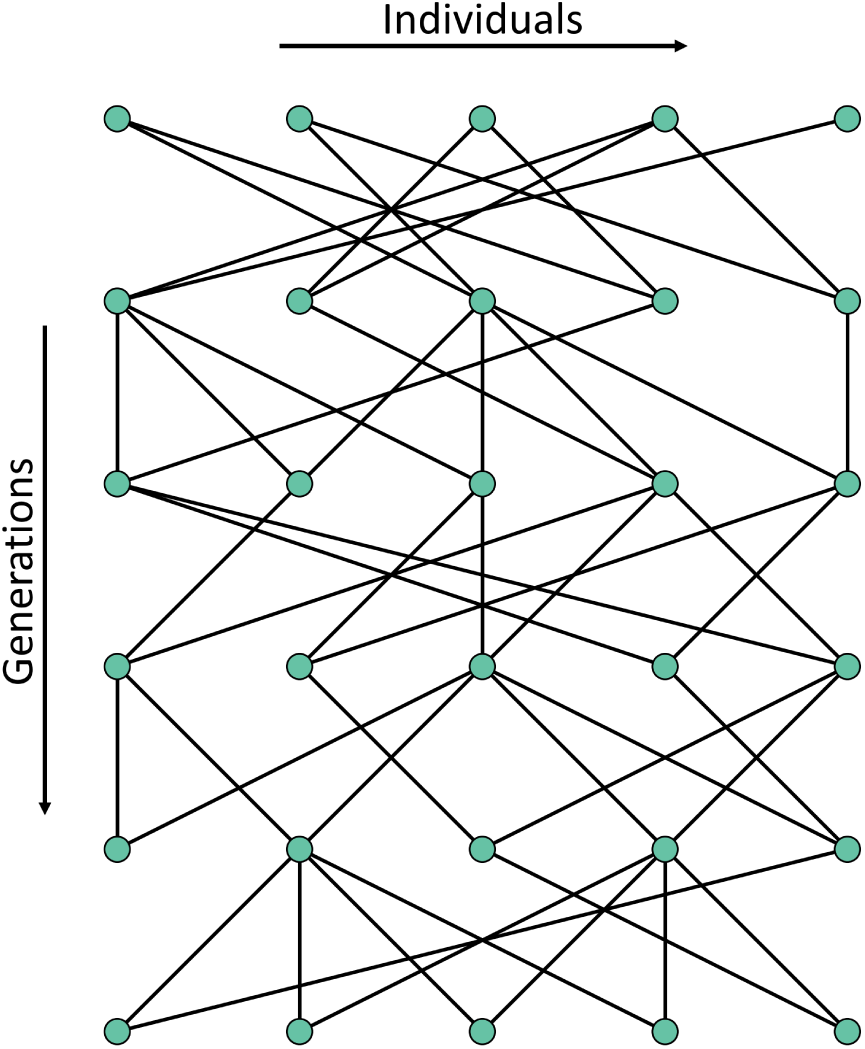
Example of pedigree in a population with biparental inheritance. Pedigree of a population resembling a Wright-Fisher population with discrete generations. Notably, each individual is assigned two random parents from the previous generation, contrary to the typical single parent inheritance in a standard Wright-Fisher model.

Recombination positions are determined by first sampling a binomial distribution to establish the number of recombination events (Figure 2), followed by sampling a uniform distribution to determine their positions. Independent assortment break positions are determined by sampling a binary random variable to ensure that each chromosome is inherited independently of the others.

**Figure 2:**
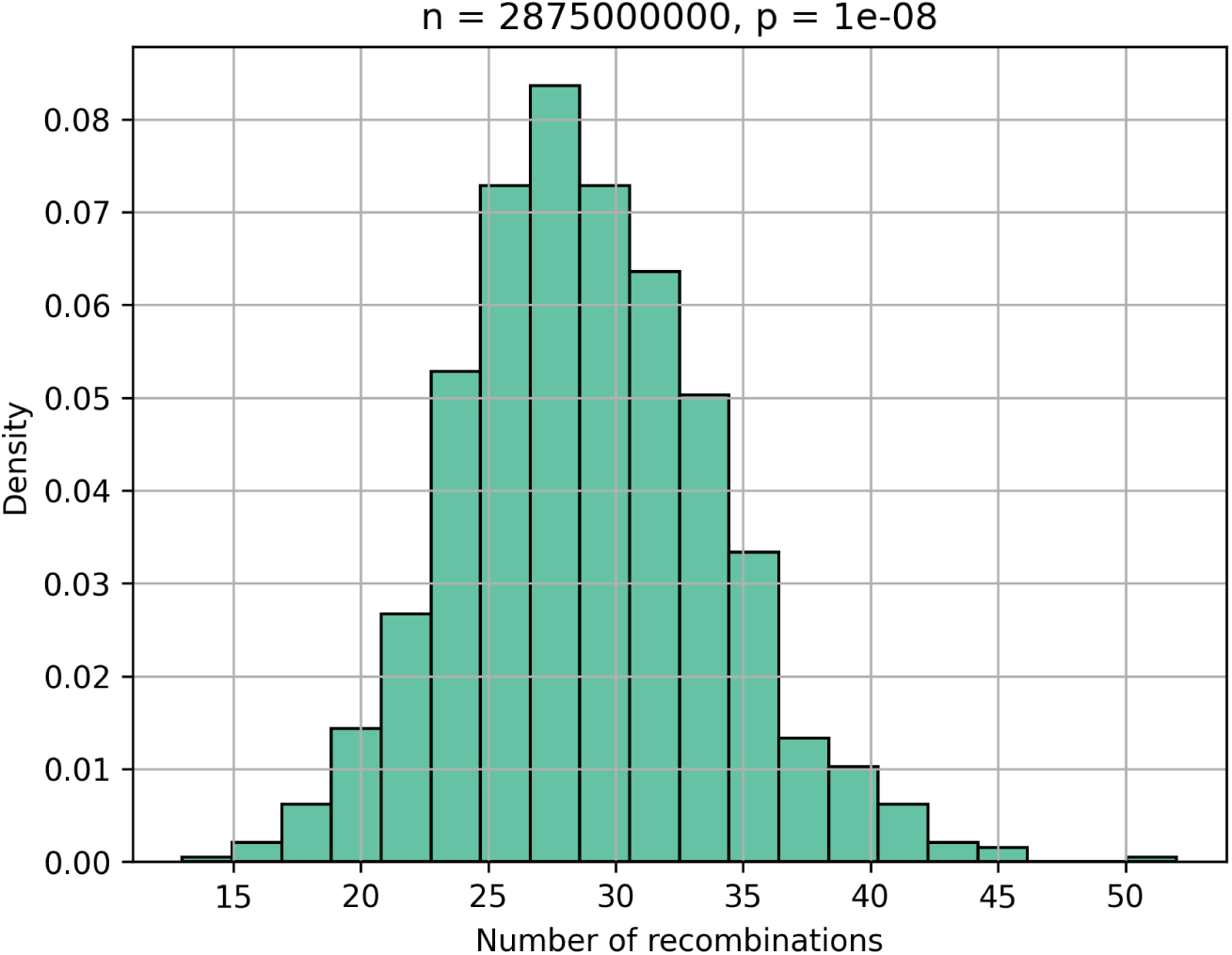
Binomial distribution representing the number of recombination events. The number of trials is 2875 *·* 10^6^ equal to the sequence length of the 22 autosomes (19). The success probability for each trial is the recombination rate of 10*^−^*^8^. The histogram shows that the number of recombination events closely resembles Coop’s assumption of 33 events (15).

### Find genealogical relatives across individuals

To identify genealogical ancestors across individuals, the model employs the breadth-first search (BFS) algorithm on a directed graph representing the population (Figure 3). All edges are directed backward in time, denoting relationships from children to parents. The BFS algorithm traverses nodes in the order of their distance from the start node and only visits each node once (20), enabling the determination of the number of ancestors per generation for a given individual.

**Figure 3:**
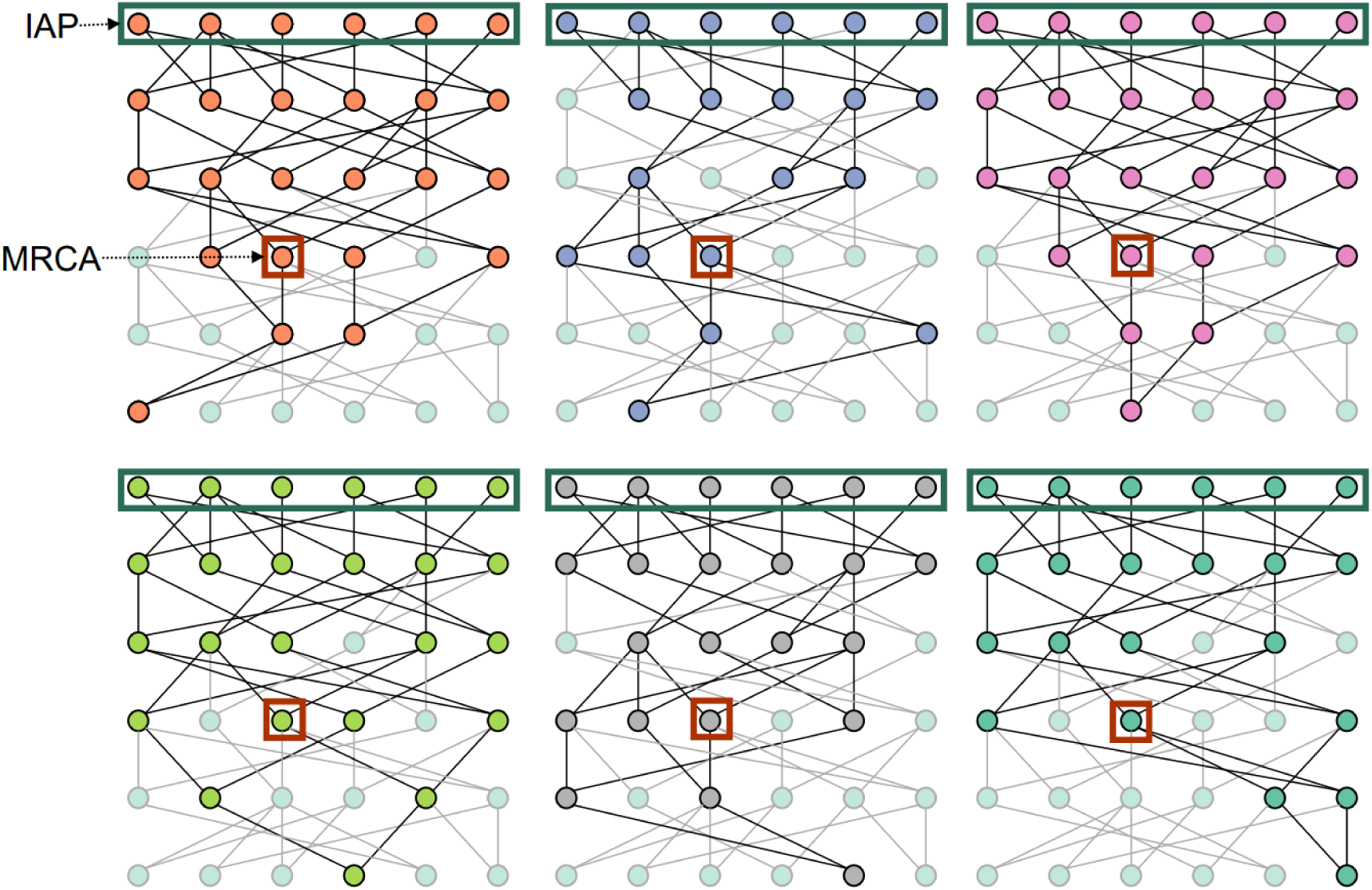
Breadth-first search (BFS) algorithm for genealogical ancestor count. Illustrating a population of 6 individuals across 6 generations, BFS is applied to each individual in the latest generation to trace genealogical ancestry. An individual’s ancestors are determined by the distinct counts in each generation. The most recent common ancestor (MRCA) is the first individual visited in all BFS traversals, marking the time of MRCA (TMRCA). The identical ancestral point (IAP) is when all individuals in a generation are ancestors, representing the point when everyone is an ancestor.

The procedure involves executing the BFS algorithm for each individual within the most recent generation and counting how many times each node has been visited in total across all BFS iterations. The time of the most recent common ancestor (TMRCA) is identified when the visit counter of one or more individuals equals the population size. The identical ancestors point (IAP) is determined when the visit counters of all individuals in a generation are either equal to the population size or zero, indicating lineage extinction.

For genealogical descendants, the same approach is employed with forward-in-time directed edges, denoting relationships from parents to children. It’s important to note that TMRCA and IAP are not applicable in descendant tracking.

### Find genetic relatedness across time

Instead of representing sequences with nucleotides, the model utilizes IDs to indicate the origin of the sequences. Each sequence is a set of ordered blocks, each assigned an ID and length. Each individual in the first generation receives a pair of distinctive blocks corresponding to their index within the generation. Individuals in the subsequent generations inherit blocks from both parents, with break positions determining transitions between sequences found within a single parent. To facilitate this process, the model implements a function to concatenate a new sequence together, forming a mosaic of the genetic material sourced from a single parent (Figure 4).

**Figure 4:**
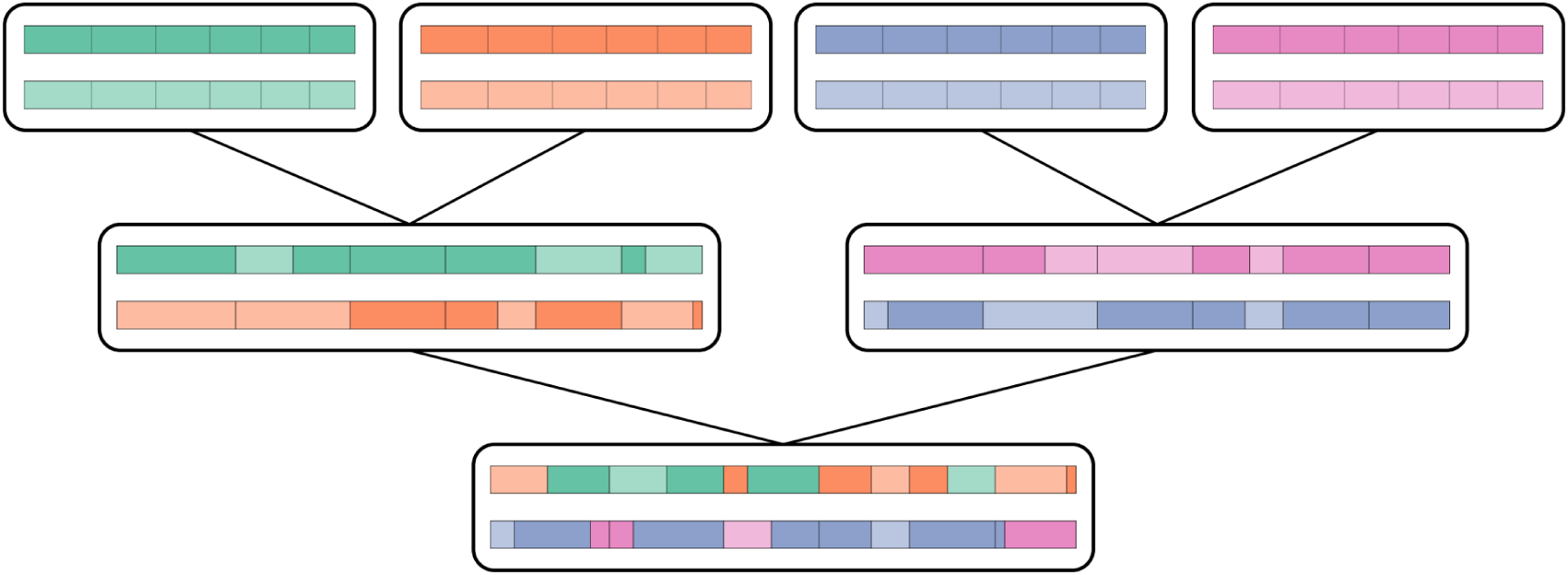
Illustration of gene spread across three generations. The process of gene spread spanning three generations, showcasing the distinct genetic contributions from grandparents through color representation. As the genetic material undergoes meiosis and recombination, observable changes occur, leading to a mosaic of genetic material with diverse origins in subsequent generations.

The number of descendants of each individual in the earliest generation is computed by counting distinct blocks within individuals in subsequent generations (Figure 5). Furthermore, the model computes the average block length across generations. The model calculates the lengths and the frequency of ROH by matching block IDs along the sequences within an individual (Figure 6) using the following equations:

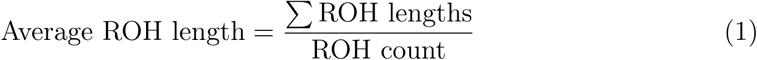

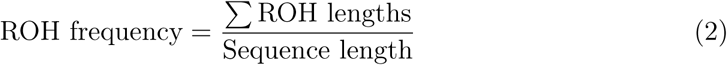

**Figure 5:**
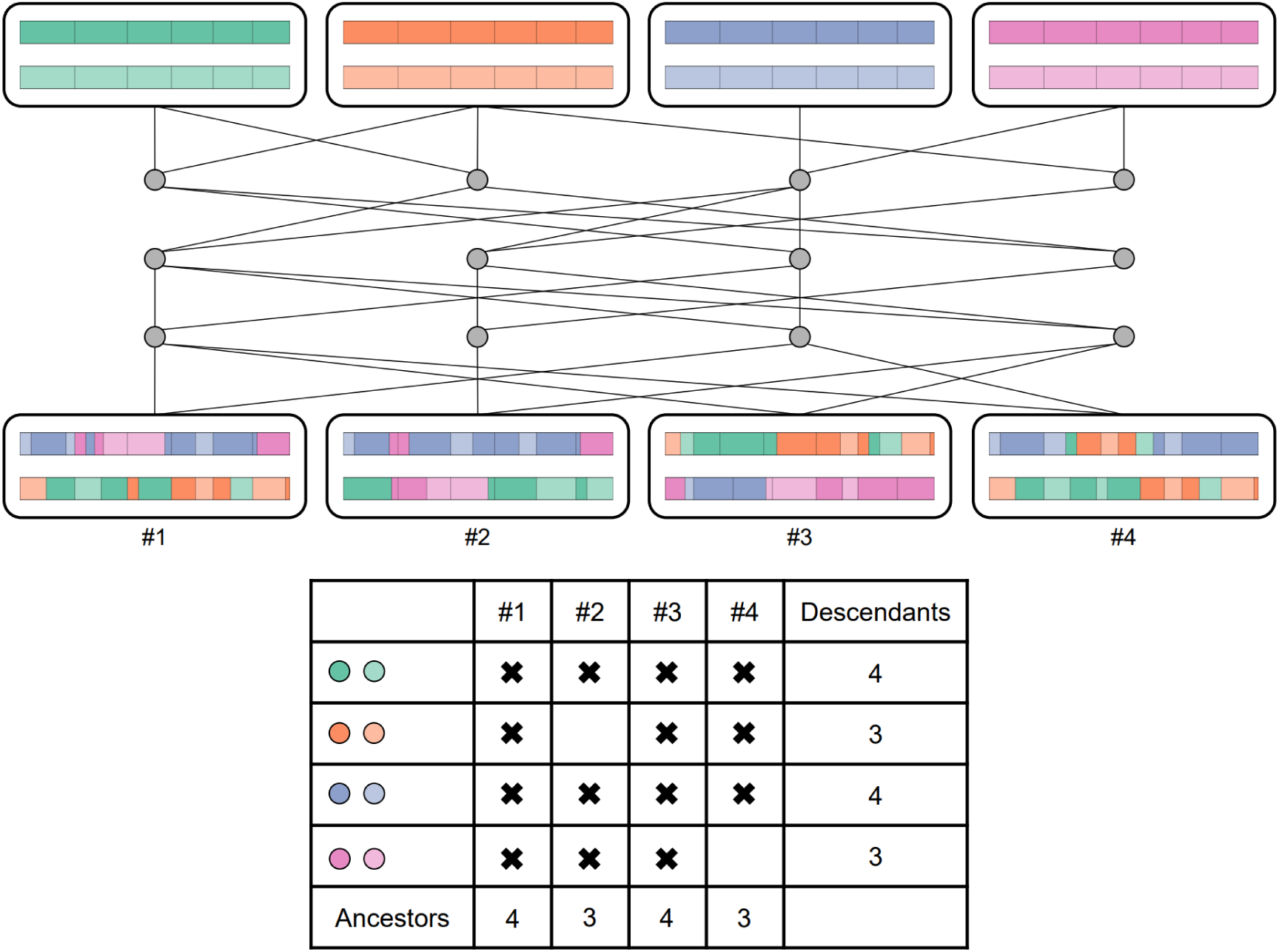
Example of count of genetic relatives across time. The upper part of the figure represents gene spread spanning multiple generations. In the lower part, a counting matrix is shown, with columns representing individuals from the most recent generation and rows representing individuals from the origin generation. Each discrete haplotype block’s color represents the genetic contribution from the origin generation to the corresponding individual in the most recent generation. The sum of each row indicates the total number of ancestors of the individual in the origin generation, while the sum of each column represents the number of descendants of the corresponding individual in the most recent generation.

**Figure 6:**
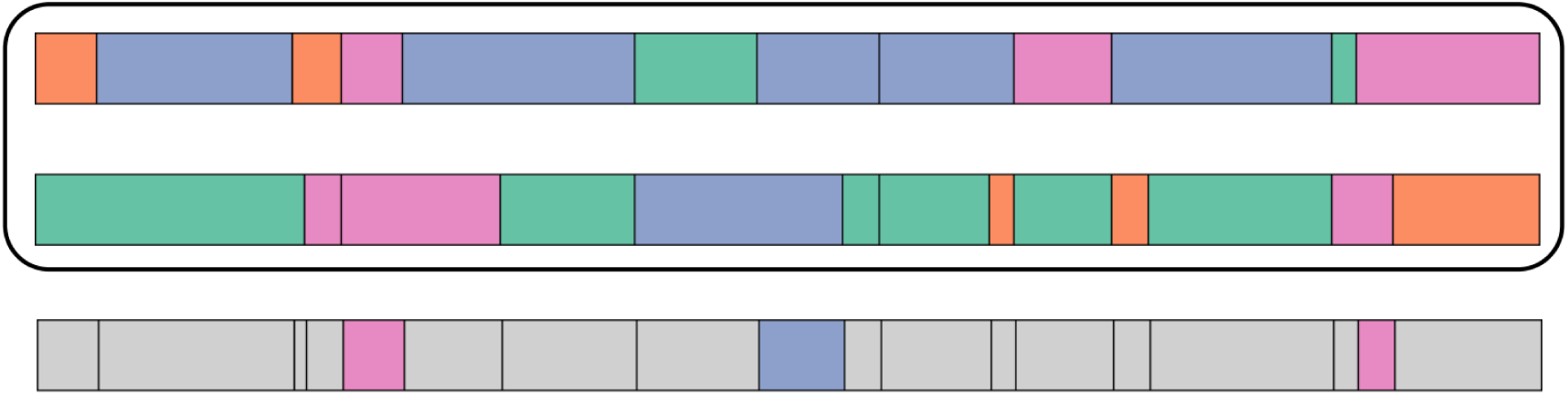
Example of runs of homozygisity (ROH) calculations. The concept of ROH illustrated through an example of two homologous sequences within an individual. Below, a sequence with break positions outlining the blocks combined from both sequences. ROH blocks, characterized by identical origins in both sequences, are highlighted. These blocks signify stretches of the genome where the individual is homozygous due to identical alleles inherited from both parents.

Counting ancestors is more complex than tallying descendants since tracing genetic material backward may result in overlapping blocks. A feature of the method is that the number of distinct blocks in the sequences of an individual corresponds to the number of ancestors in the earliest generation of that individual (Figure 5). To determine the number of ancestors over generations, the analysis is repeated for the same population, starting from different generations. Analyzing one generation back, then two, and so on, up to a set number of generations. In each iteration, distinct blocks in the sequences of individuals from the latest generation are counted. Adding up these counts gives the average number of ancestors across generations.

### Utilizing the proposed model

We used the proposed model to quantify the expected numbers of ancestors and descendants, the counts and lengths of haplotype blocks, and lengths and frequency of ROH. This involved running the model across various population scenarios. The simulation parameters included population sizes of 50, 100, 250, 500, and 1000 individuals, spanning 100 generations. The recombination rate was set at 10*^−^*^8^ with a random seed value of 395. Chromosome lengths were obtained from Piovesan *et al.* (19).

Furthermore, to analyze the distribution of ROH frequency, we performed 10 simulations for 50 generations, again for various population sizes of 50, 100, 250, 500, and 1000 individuals. Additionally, we ran a simulation with a population size of 50 for 1000 generations to test long-term metrics. To assess the impact of population size, we extended our simulations to larger populations of 10,000, 25,000, and 100,000 individuals, allowing us to observe trends over time before the effects of population size become significant.

## Results

### Number of ancestors

Our analysis of the number of ancestors delves into population dynamics across varying sizes and generations. Initial exploration focuses on early trends, examining populations of 50 and 100 individuals over the initial 25 generations (Figure 7). Notably, both genetic and genealogical ancestors exhibit an exponential growth pattern in preliminary generations. However, this rapid growth is curtailed, stabilizing at approximately 80% of the population size. Furthermore, we extend our investigation by exploring population sizes of 250, 500, and 1000 individuals, spanning 100 generations (Figure 8). The time of most recent common ancestor (TMRCA) and the identical ancestors point (IAP) are marked in both figures. Our findings indicate that the exponential growth phase extends over a longer duration for larger population sizes.

**Figure 7:**
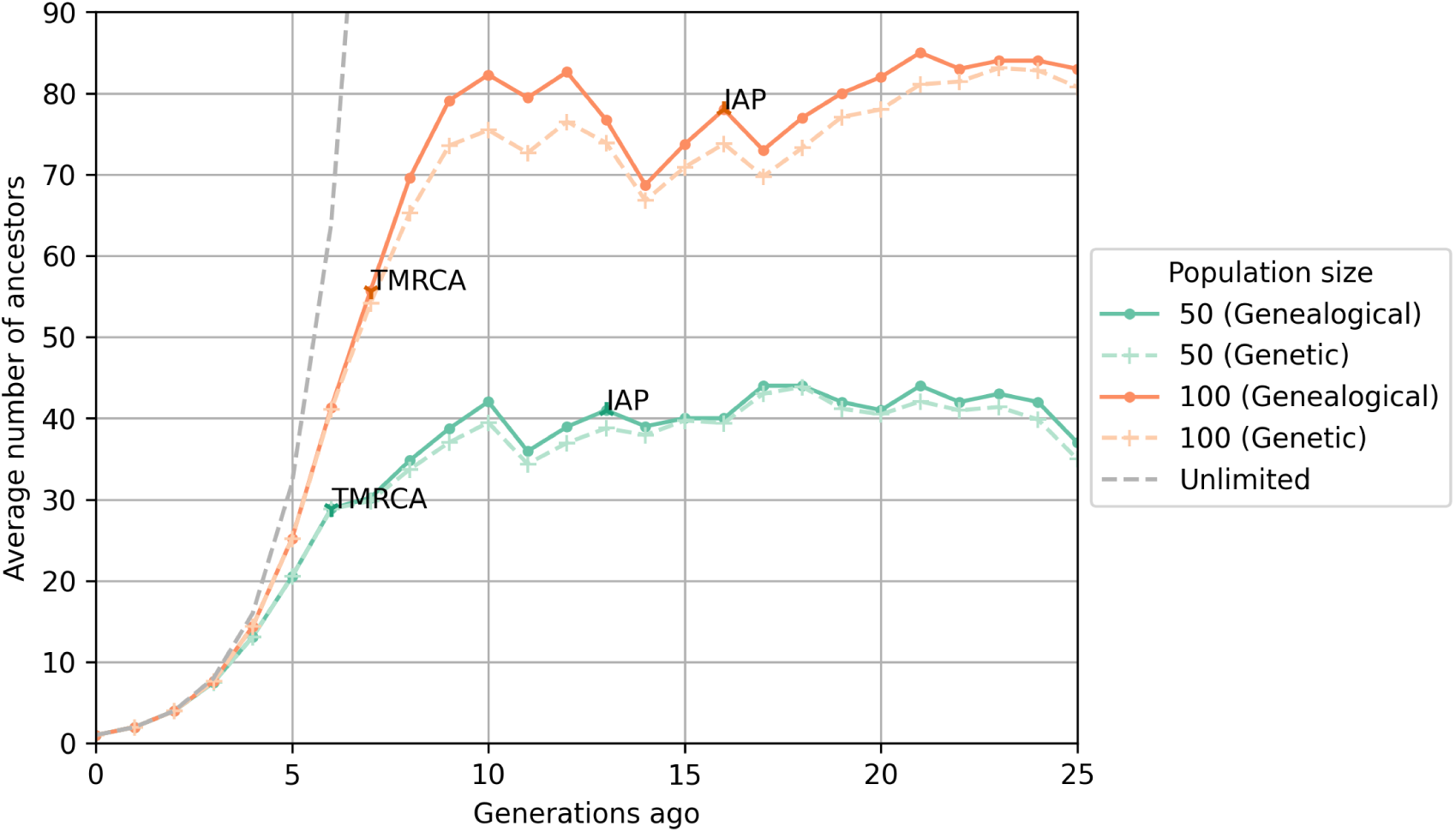
Number of ancestors in initial generations. Average number of genetic and genealogical ancestors for population sizes of 50 and 100 individuals over 25 generations. The figure also shows the exponential growth for a population of unlimited size.

**Figure 8:**
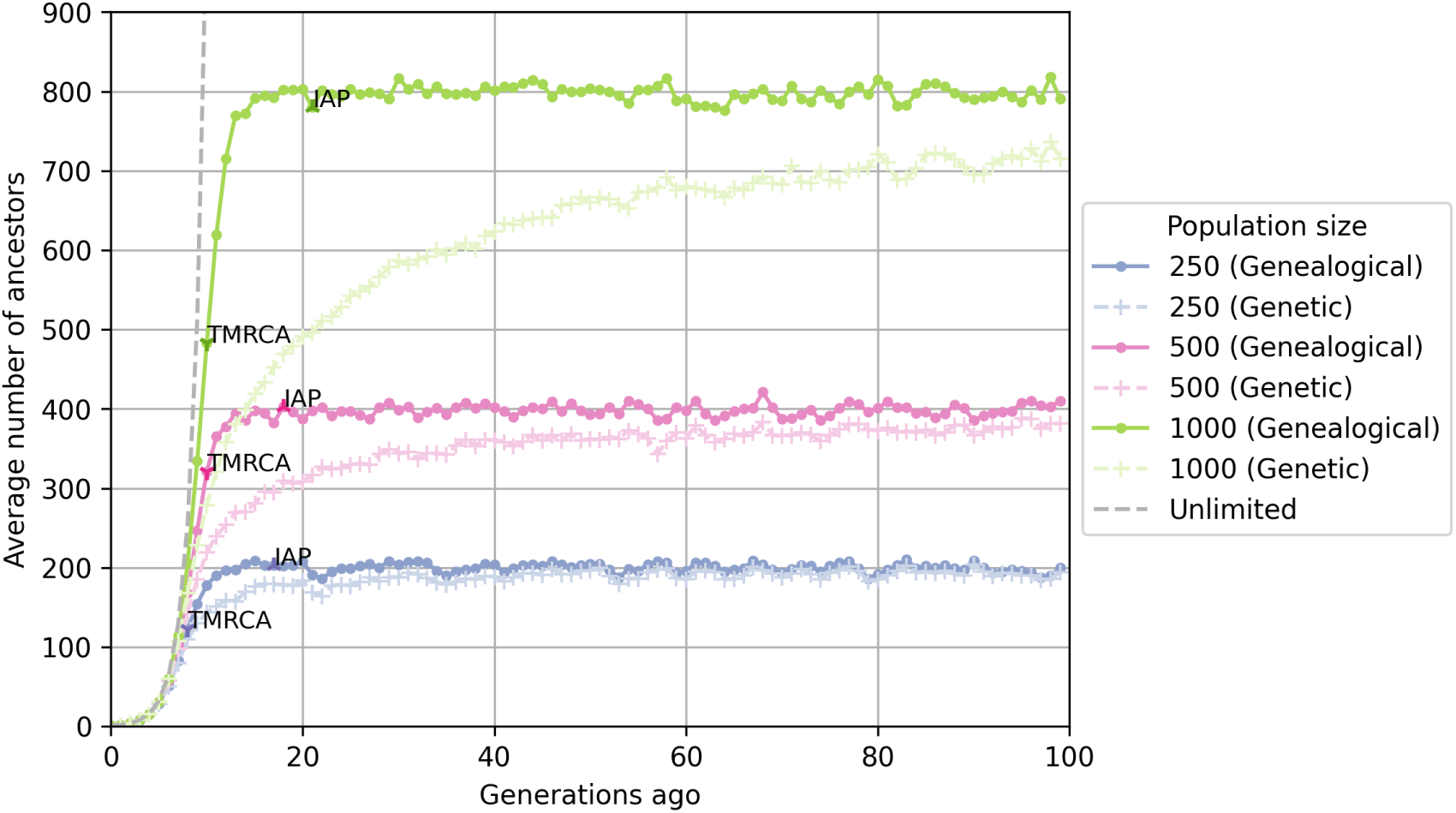
Number of ancestors in long-term analysis. Average number of genetic and genealogical ancestors for population sizes of 250, 500, and 1000 individuals over 100 generations. The figure also shows the exponential growth for a population of unlimited size.

Additionally, we compare our results with those from Coop (15). Simulations for much larger population sizes of 10,000, 25,000, and 100,000 individuals across 25 generations reveal similar trends in both datasets, although our simulations yield lower ancestor counts (Figure 9).

**Figure 9:**
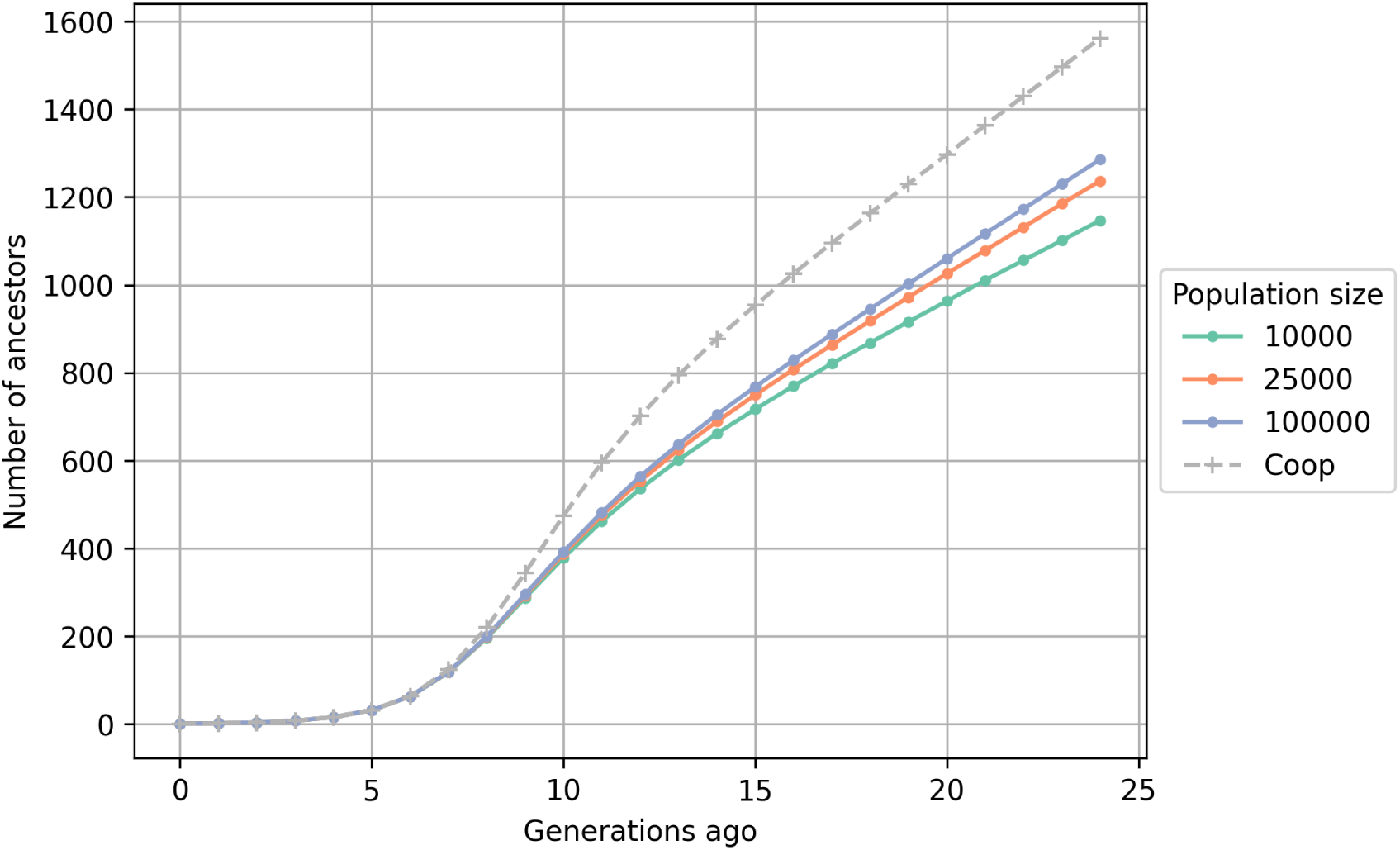
Number of ancestors for large populations. Average number of genetic ancestors for population sizes of 10,000, 25,000, and 100,000 over 25 generations in comparison with Coop’s results (15).

### Number of descendants

Extending our investigation to the average number of descendants per subsequent generation, we consider both early trends and long-term patterns. Figures 10 and 11 present the average number of descendants of individuals in the earliest generation who had offspring.

**Figure 10:**
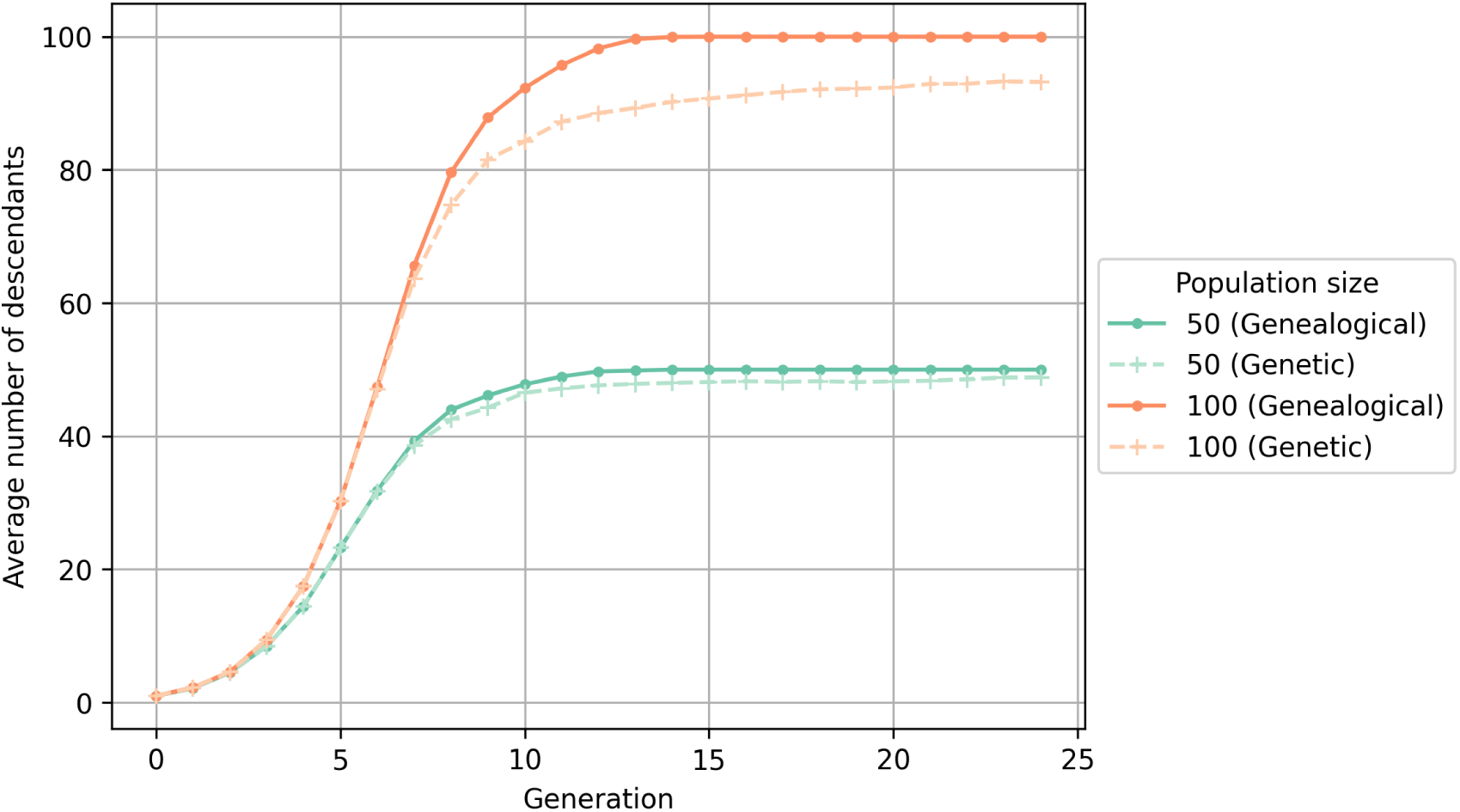
Number of descendants in initial generations. Average number of genetic and genealogical descendants for population sizes of 50 and 100 individuals over 25 generations.

**Figure 11:**
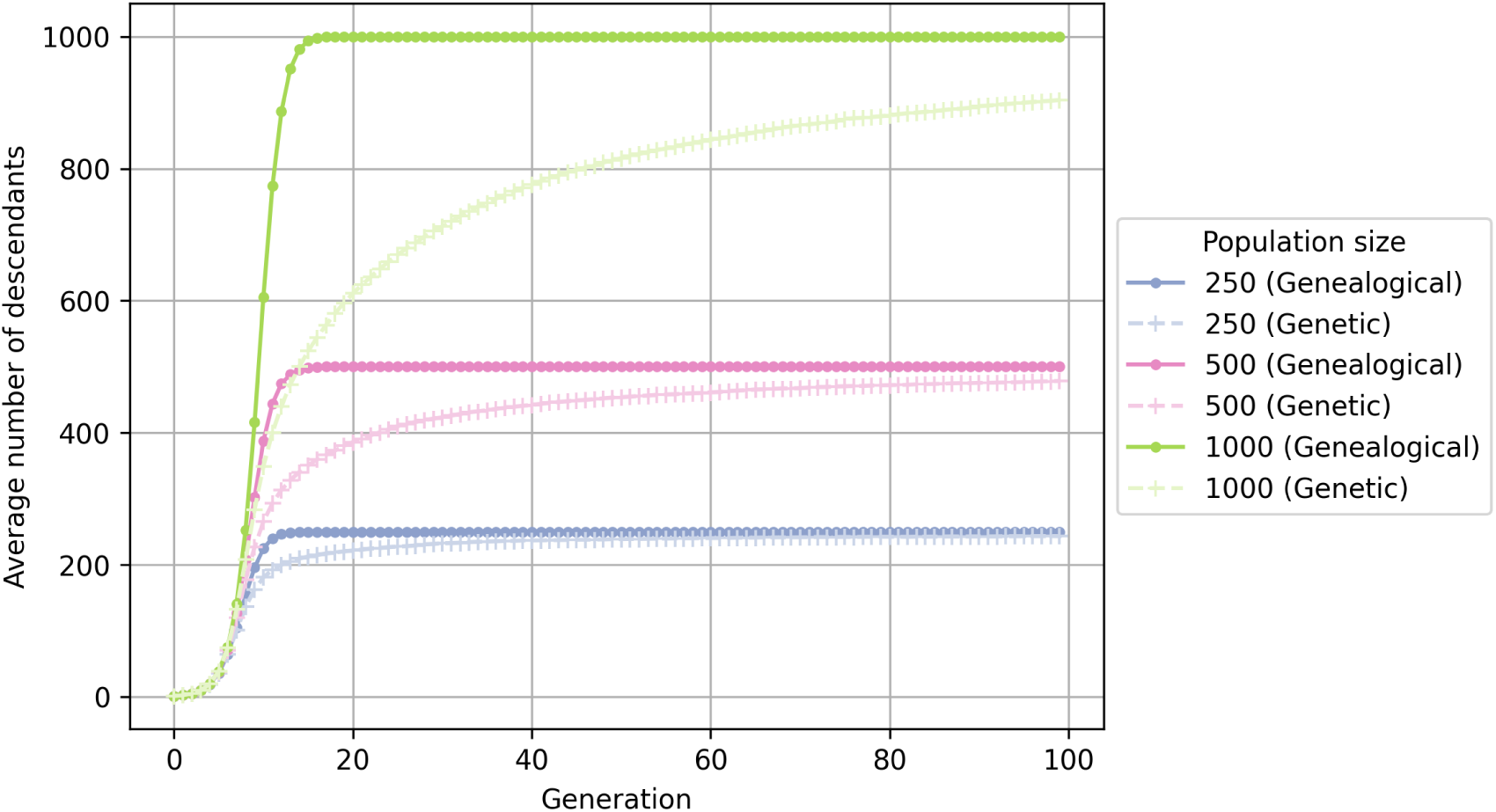
Number of descendants in long-term analysis. Average number of genetic and genealogical descendants for population sizes of 250, 500, and 1000 individuals over 100 generations.

Consistent with ancestral findings, the initial generations exhibit an exponential growth phase in the number of descendants, eventually converging against the population size. Unlike the ancestral data, the number of descendants remains more stable, showing less fluctuation.

### Haplotype blocks and runs of homozygosity

In this section, we explore the analysis of both the length of haplotype blocks and the length and frequency of runs of homozygosity (ROH) across time. Our investigation into the length of haplotype blocks across time (Figure 12) indicates a consistent decrease over generations, with no significant influence from changes in population size.

**Figure 12:**
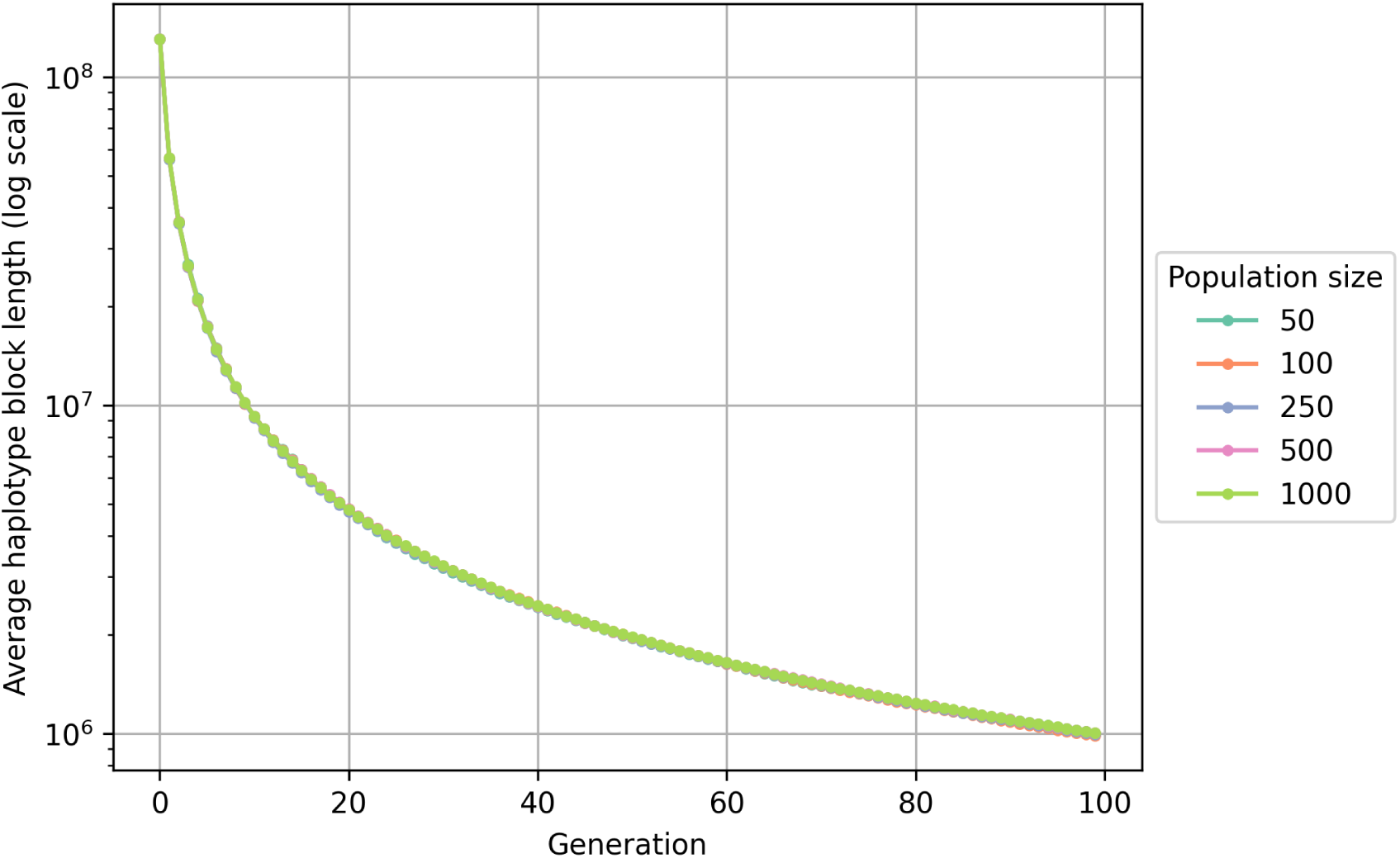
Length of haplotype blocks across time. Haplotype block lengths are depicted for population sizes of 50, 100, 250, 500, and 1000 individuals across 100 generations. The y-axis is presented on a logarithmic scale to better visualize the data trends.

The length of ROH segments (Figure 13) appears to be unaffected by changes in population size, similar to what we observed with haplotype blocks. However, the expected frequency of ROH (Figure 14) increases over time, with higher values associated with smaller population sizes. To showcase the long-term trend of ROH, we include results for a population of 50 individuals across 1000 generations (Figure 15). In this scenario, we observe that after approximately 700 generations, the ROH reaches 100%.

**Figure 13:**
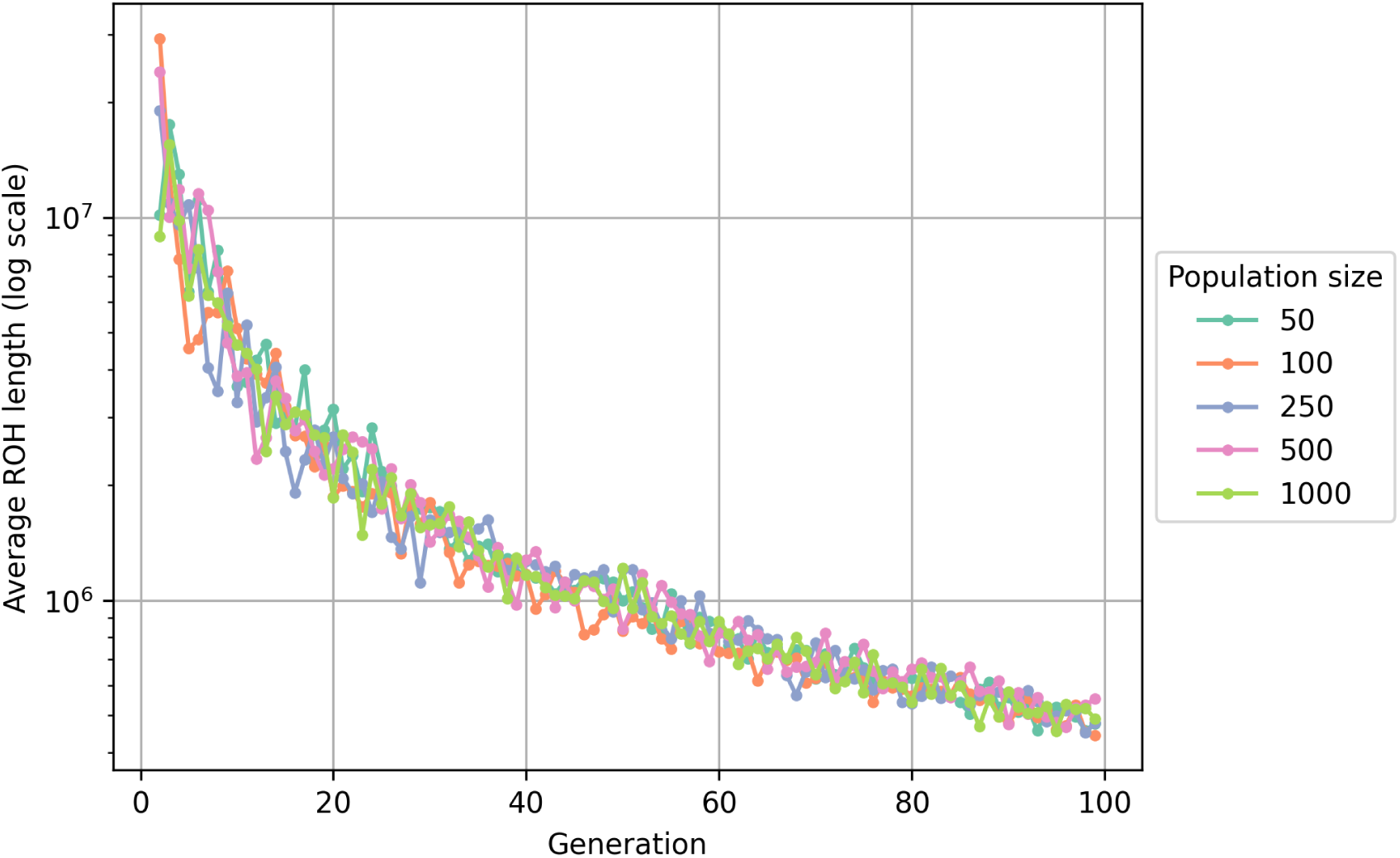
Length of ROH across time. ROH lengths are depicted for population sizes of 50, 100, 250, 500, and 1000 individuals across 100 generations. The y-axis is presented on a logarithmic scale to better visualize the data trends.

**Figure 14:**
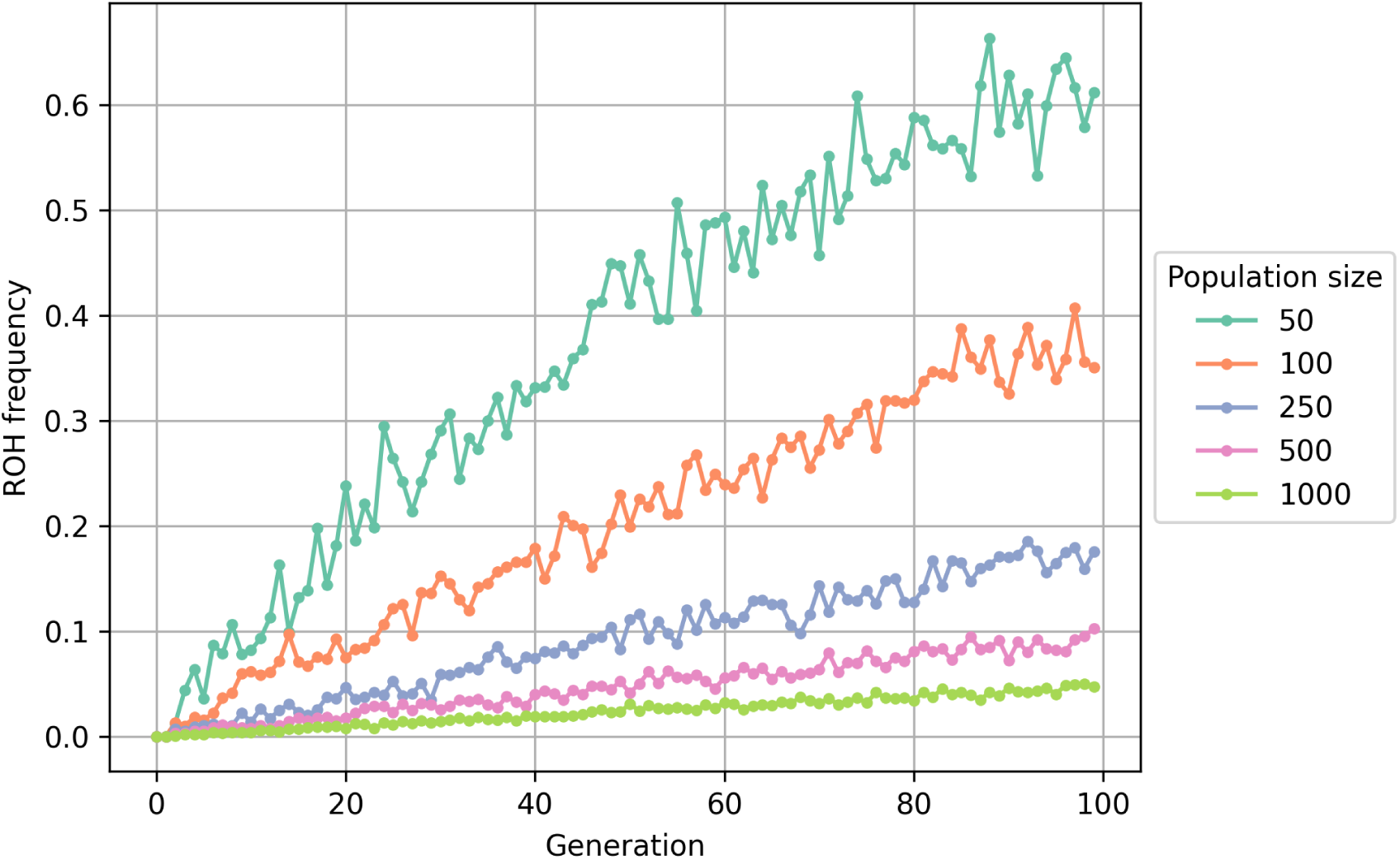
Frequency of ROH across time. Distribution of the frequency of ROH for population sizes of 50, 100, 250, 500, and 1000 individuals over 100 generations.

**Figure 15:**
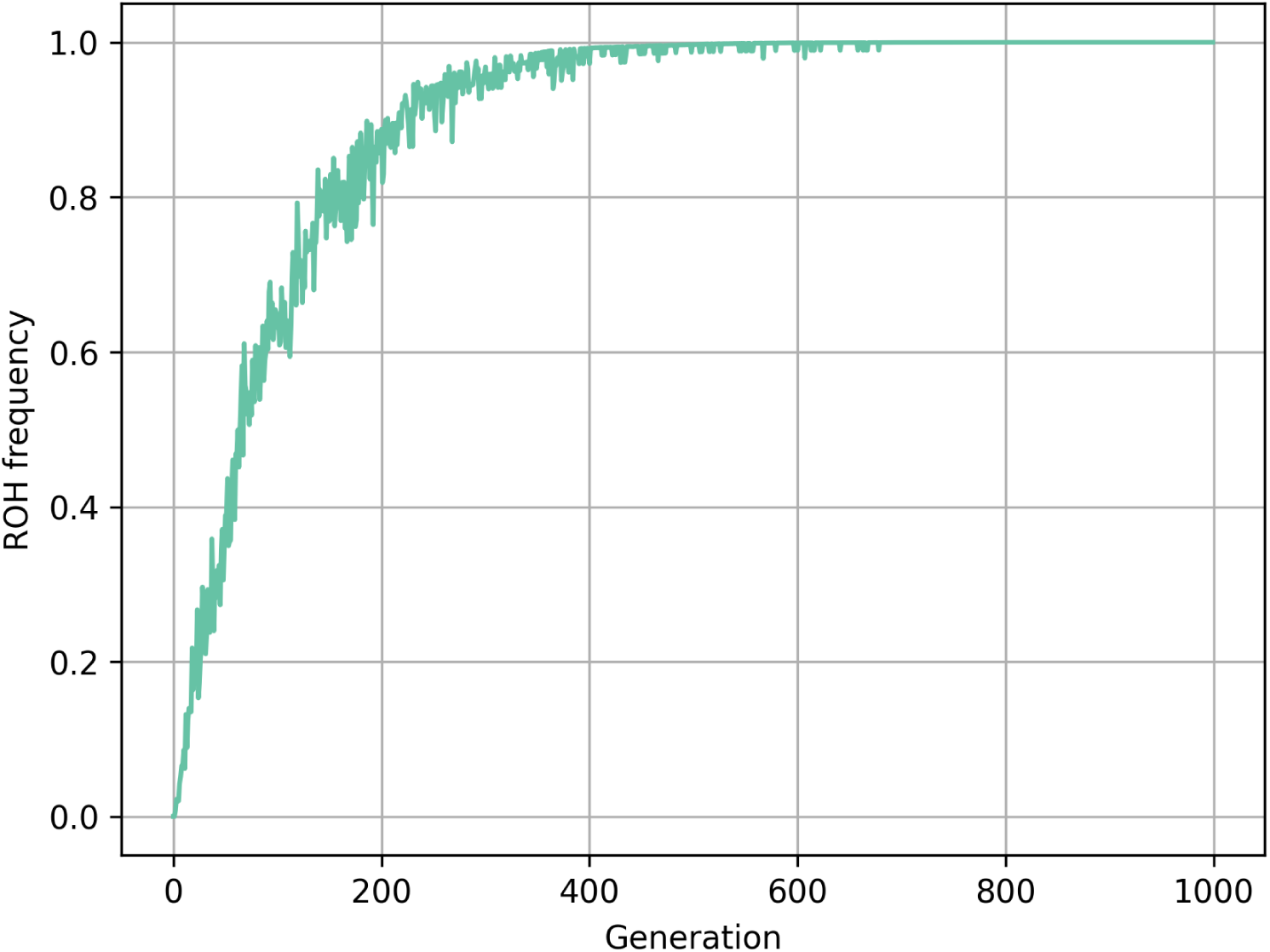
Long-term trend of ROH over 1000 generations. Analysis of the frequency of ROH over 1000 generations for a population size of 50 individuals.

In order to assess the variability in the frequency of ROH across different population scenarios, we conducted a series of simulations. Each simulation spans 50 generations, with 10 runs for each population size. Figure 16 presents a boxplot illustrating the distribution of ROH frequency along with its deviation for each population scenario. Notably, for a population size of 50 individuals, we observe a ROH frequency averaging just under 40% across all simulations. Conversely, when the population size increases to 1000 individuals, the ROH frequency decreases significantly, stabilizing at approximately 2.5% on average.

**Figure 16:**
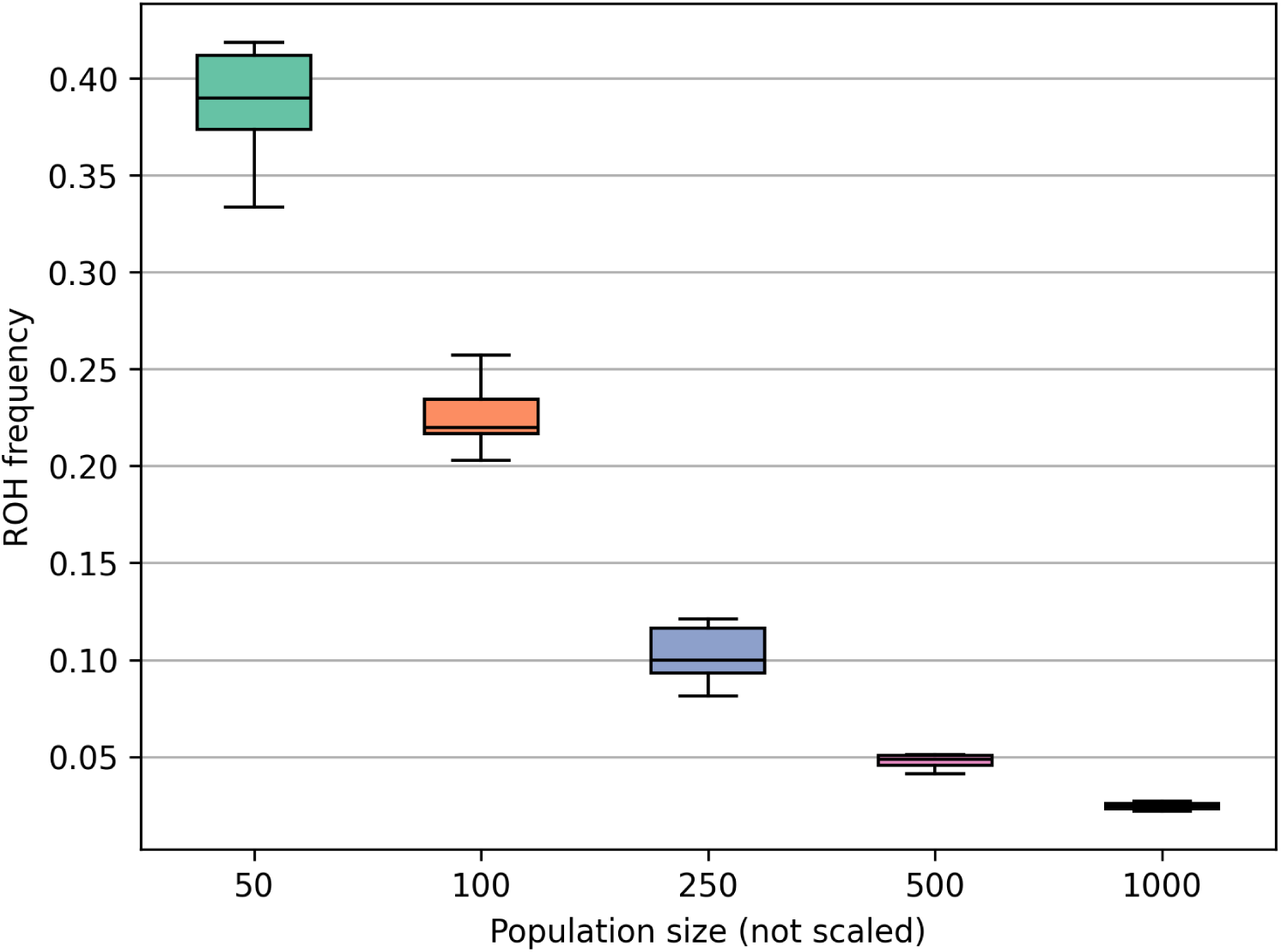
Variability in ROH frequency across simulations for different population scenarios. Variability in the frequency of ROH across simulations for population sizes of 50, 100, 250, 500, and 1000 individuals. The x-axis is unscaled.

## Discussion

### Genealogical and genetic ancestors

Our analysis of the number of ancestors as we trace back in time provides insights into the interplay between population size and genetic heritage. One observation is the initial exponential growth in the number of ancestors, a consequence of each individual having two parents, four grandparents, and so on. However, this exponential growth is constrained by the finite size of the population, leading to a transition into logistic growth.

The duration of the exponential growth phase is influenced by population size, with larger populations exhibiting longer periods of exponential growth. Still, even in smaller populations, the exponential phase is relatively brief due to the limiting effect of population size. This observation aligns with our expectations, as in a small population, the genetic diversity is rapidly exhausted, leading to a convergence towards a common set of ancestors.

In contrast to genealogical ancestors, genetic ancestors exhibit a more variable growth pattern across different population sizes. The influence of recombination and independent assortment becomes apparent here, allowing genes from different ancestors to be passed on to offspring. Consequently, the number of genetic ancestors continues to grow over many generations, in contrast to the stabilization observed in genealogical ancestry.

Notably, the number of ancestors converges to around 80% of the population size, a point reached around the time of the most recent common ancestor (TMRCA). This convergence underscores an aspect of genetic ancestry: as we go back in time, a decreasing number of individuals are responsible for contributing genetic material to the entire population.

The fluctuations observed around this 80% point can be attributed to the stochastic nature of our model, where not all individuals have offspring in every generation, leading to variations in the number of individuals contributing genetic material. Despite these fluctuations, this percentage appears to be a reasonable estimate, supported by calculations based on the expected number of distinct draws from a finite population.

### Comparison with Coop’s results

In our analysis, we compared our results with Coop’s for the genetic number of ancestors (15; 16). Coop is using a formula, derived from probability theory, predicts a higher number of genetic ancestors and a more extended period of exponential growth. Nevertheless, some similarities between the two methods are evident, particularly in the shape of the growth curves.

The difference between the two methods can likely be attributed to the finite population size we considered in our simulations. While we did not explore larger population sizes extensively, our observations from simulations with population sizes ranging from 10,000 to 100,000 individuals suggest that the differences between the methods reduce with larger populations. This supports the notion that our finite population size has a noticeable effect on the outcomes and highlights the importance of considering population size when analyzing genetic ancestry.

### Genetic descendants

Our analysis of genetic descendants provides insights into the expected relatedness of individuals moving forward in time. The trend of logistic growth observed mirrors that of genetic ancestors, with smaller populations converging faster, indicating closer genetic relatedness in smaller populations. The stability observed in this convergence, as compared to when we look backward in time, can be attributed to the fact that while every individual has parents, not everyone becomes a parent, and that is why we don’t see the fluctuations in the numbers here.

The effect of meiosis becomes evident in the difference between the number of genetic and genealogical ancestors. Meiosis results in the halving of an individual’s genetic material passed to offspring, contributing to the gap between genetic and genealogical relatedness. This observation underscores the significance of genetic processes in shaping the relatedness of individuals over time.

### Haplotype blocks

Turning our attention to the length of haplotype blocks, we observed a consistent decrease in the length of haplotype blocks, unaffected by population structure. This phenomenon can be attributed to the effect of genetic recombination, where crossovers between chromosomes during meiosis result in the shuffling of genetic material between homologous chromosomes. This observation aligns with our expectations and underscores the dynamic nature of genetic inheritance.

### Runs of homozygosity

Our analysis of ROH lengths reveals a trend that closely resembles the length distribution of haplotype blocks. This observation suggests that, like haplotype blocks, the population structure did not significantly influence the distribution of ROH lengths. However, there was a notable difference in the stability of these lengths. While haplotype block lengths decreased in a relatively stable manner, we observed fluctuations in ROH lengths. This phenomenon could be attributed to the fact that ROH segments are generally fewer in count compared to haplotype blocks, and the limited number of ROH segments could lead to fluctuations in their lengths. However, when we considered a high-frequency threshold of 80% (Figure 15), indicating that a substantial portion of the population shared these segments, we still observed a high degree of fluctuations.

The ROH analysis also shed light on the significant impact of population size on the expected frequency of ROH. Smaller populations showed a much higher degree of ROH, consistent with the principles of genetic drift, where in smaller populations, random sampling of alleles during reproduction can lead to a greater chance of homozygosity. Similar to the patterns observed in ROH segment lengths, we also noted fluctuations in ROH frequency within individuals.

To illustrate our findings, we presented Figure 16, which shows the results of the average ROH frequency after 50 generations for various population sizes. This figure provides a clear visualization of the mean expected frequency of ROH and their distribution. We see that for a small population of 50 individuals, we can expect almost 40% ROH after 50 generations. In contrast, when the population size increases to 1000 individuals, the expected ROH frequency decreases significantly to 2.5%.

Notably, our model did not account for the evolutionary force of mutation. While this omission did not significantly affect our analysis of the number of relatives, it likely overestimated the frequency of ROH. The introduction of mutations could lead to increased genetic diversity, potentially altering the ROH patterns observed in our simulations. Future iterations of the model could consider incorporating mutation rates to refine ROH predictions.

### Understanding ROH patterns in drifted populations

Our analysis of expected ROH frequency offers valuable insights into the degree of ROH within highly drifted populations. This insight is particularly relevant when considering populations with a history characterized by founder effects, bottlenecks, and limited gene flow, such as the Faroe Islands.

The Faroe Islands serve as a compelling example of a highly drifted population. Founded by a relatively small number of individuals approximately 50 generations ago, this population experienced low population growth rates until the 1800s and minimal migration or gene flow with other populations. Additionally, historical events, such as the plague outbreak in the 1300s (8), further reduced the effective population size due to a bottleneck effect.

Despite the Faroe Islands currently having a population size exceeding 50,000 individuals, the effective population size is believed to be significantly smaller, possibly even lower than 1,000 individuals. This substantial reduction in effective population size has had a profound impact on the genetic composition of the population.

In this context, Figure 16 becomes a valuable tool for gaining insights into the expected ROH frequency in populations like the Faroe Islands. By extrapolating from our analysis, we can make informed estimations about the prevalence of ROH in highly drifted populations, shedding light on the genetic consequences of their unique demographic histories.

This perspective allows us to appreciate how our research not only contributes to our understanding of general population genetics principles but also provides practical applications for assessing and interpreting the genetic makeup of populations with complex histories of isolation and demographic change, like the Faroe Islands.

## Conclusion

In conclusion, our analysis of genetic and genealogical ancestors, genetic descendants, haplotype blocks, and runs of homozygosity (ROH) has provided valuable insights into the complex dynamics of genetic inheritance, population size, and the effects of genetic processes.

We found that the relationship between population size and genetic heritage shapes the number of ancestors, with exponential growth transitioning into logistic growth due to finite population constraints. Genetic ancestors, influenced by recombination and independent assortment, exhibit a more variable growth pattern but eventually converge to around 80% of the population size.

Comparing our results with Coop’s genetic ancestor model highlighted the significance of population size in genetic ancestry analysis. Our findings suggest that finite population size can lead to differences in outcomes, emphasizing the importance of considering population size when studying genetic ancestry.

The analysis of genetic descendants revealed trends in relatedness, with smaller populations showing faster convergence. Meiosis and recombination played a critical role in the gap between genetic and genealogical relatedness due to halving an individual’s genetic material.

Analyzing ROH, we found that ROH lengths closely resembled haplotype block length distributions. Population size significantly influenced ROH frequency, with smaller populations exhibiting higher ROH levels due to genetic drift. This insight is particularly relevant when considering highly drifted populations like the Faroe Islands, where ROH patterns can shed light on the genetic consequences of their unique demographic histories.

Our findings indicate that ROH lengths closely resemble haplotype block length distributions, with a significant impact of population size on ROH frequency. For instance, in a small population of 50 individuals, we can expect almost 40% ROH after 50 generations. In contrast, when the population size increases to 1000 individuals, the expected ROH frequency decreases significantly to 2.5%. These numerical results illustrate the pronounced effect of population size on ROH levels, aligning with principles of genetic drift.

By developing a comprehensive population simulation model, we have created a tool for exploring the intricate relationships between genetic inheritance, demographic changes, and the effects of different evolutionary forces in various population scenarios.

## Acknowledgements

We would like to acknowledge that Gabriel Renaud is funded by the Novo Nordisk Data Science Investigator grant number NNF20OC0062491. We would like to thank Peter Wad Sackett and the Department of Healthtech at the Technical University of Denmark (DTU).

